# SAM-AMP lyases in CRISPR defence and anti-defence

**DOI:** 10.1101/2025.04.28.651013

**Authors:** Haotian Chi, Stephen McMahon, Lukas Daniel-Pedersen, Shirley Graham, Tracey M Gloster, Malcolm F White

## Abstract

Type III CRISPR systems detect non-self RNA and activate the enzymatic Cas10 subunit, which generates nucleotide second messengers for activation of ancillary effectors. Although most signal via cyclic oligoadenylate (cOA), an alternative class of signalling molecule SAM-AMP, formed by conjugating ATP and S-adenosyl methionine, was described recently. SAM-AMP activates a trans-membrane effector of the CorA magnesium transporter family to provide anti-phage defence. Intriguingly, immunity also requires SAM-AMP degradation by means of a specialised CRISPR-encoded NrN family phosphodiesterase in *Bacteroides fragilis*. In *Clostridium botulinum*, the *nrn* gene is replaced by a gene encoding a SAM-AMP lyase. Here, we investigate the structure and activity of *C. botulinum* SAM-AMP lyase, which can substitute for the *nrn* gene to provide CorA-mediated immunity in *Escherichia coli*. The structure of SAM-AMP lyase bound to its reaction product methylthioadenosine-AMP (MTA-AMP) reveals key details of substrate binding and turnover by this PII superfamily protein. Bioinformatic analyses reveal candidate phage-encoded SAM-AMP lyases and we demonstrate that one, hereafter named AcrIIIB4, degrades SAM-AMP efficiently *in vitro*.

## INTRODUCTION

CRISPR-Cas systems function as an adaptive immune system in bacteria and archaea against mobile genetic elements (MGE) such as viruses and plasmids. Type III systems bind RNA from MGE, resulting in the activation of the enzymatic Cas10 subunit, which typically comprises a specialised polymerase activity that generates a family of cOA signalling molecules from ATP (1,2). cOA acts as a second messenger of infection in the cell, activating a wide range of ancillary effectors to provide an immune defence. These include nucleases such as Csm6 (2), Csx1 (3), Can1 (4), Can2 (5,6) and NucC (7,8) (reviewed in (9)), proteases such as CalpL (10) and SAVED-CHAT (11), transcriptional regulators such as Csa3 (12), translational inhibitors such as Cami1 (13) and transmembrane effectors such as Csx23 and Cam1 (14,15). Recently, effector distribution has been analysed and new effector families proposed (16). Enzymes known as ring nucleases, which can be CRISPR- or virally-encoded, degrade cOA to deactivate type III CRISPR systems (17-21).

Recently, a new class of signal molecule, SAM-AMP, has been identified in a *Bacteroides fragilis* type III-B CRISPR system (22). Two accessory proteins, a CorA family transmembrane protein and an NrN-family phosphodiesterase, were shown to be essential for CRISPR-mediated immunity in the heterologous host *E. coli*. SAM-AMP binding to CorA is thought to potentiate disruption of membrane integrity, resulting in growth arrest or death of infected cells. *In vitro*, NrN functions analogously to ring nucleases, degrading SAM-AMP into SAM and AMP. While this might be expected to represent a means to switch off the immune response, plasmid challenge assays in *E. coli* suggest NrN activity is required for immunity – an unexpected observation that is still not fully understood (22). NrN is replaced by a SAM-AMP lyase in some CorA-associated type III CRISPR systems, including that from *C. botulinum* (22). Previously, phage-encoded SAM lyases have been found to degrade host SAM into 5’-methyl-thioadenosine (MTA) and L-homoserine lactone (HL), thus neutralising the bacterial restriction-modification (RM) (23) or bacterial exclusion BREX (24) systems. The phage SAM lyase exhibits structural features characteristic of PII-like signalling proteins: three ferredoxin-like folds assembled into a trimeric structure with a triangular core of β-sheets (23,25). Proteins containing this structural feature have the capability to bind ligands like ATP, c-di-AMP and MTA at the trimeric interface, enabling them to be involved in various cellular processes including the regulation of anabolic metabolism, metal homeostasis and SAM degradation (23,25).

Here, we investigate *C. botulinum* SAM-AMP lyase both *in vivo* and *in vitro*, demonstrating that it is functional in the context of *B. fragilis* CorA-associated CRISPR systems in *E. coli* and specifically degrades SAM-AMP to HL and MTA-AMP. The crystal structure of *C. botulinum* lyase bound to its MTA-AMP product provides insights into SAM-AMP recognition and turnover, as well as providing the first definitive view of a SAM-AMP derived molecule, which confirms the predicted 5’-3’ phosphodiester linkage. We go on to demonstrate that phage also encode SAM-AMP lyases, which presumably function as anti-CRISPRs (Acrs) against SAM-AMP signalling defence systems.

## MATERIALS AND METHODS

### Cloning

Cloning of the gene encoding SAM-AMP lyase from *C. botulinum* has been described previously (22). Briefly, a synthetic gene encoding SAM-AMP lyase was ordered as a g-block (IDT) and cloned between the *Nco*I and *Bam*HI restriction sites of vector pEhisV5TEV (26) for protein purification, or between the restriction sites *Nco*I and *Eco*RI of vector pRATDuet (21) or plasmid pRATDuet-BfrCorA containing *cora* between restriction sites *Nde*I and *Xho*I for plasmid challenge assays. Mutations were introduced by site-directed mutagenesis (Phusion High-Fidelity DNA polymerase, Fisher Scientific) using primers listed in Supplementary Table1.

### Expression and purification of SAM-AMP lyase

The pEhisV5TEV expression plasmid was transformed into *E. coli* C43 (DE3) cells and grown at 16 °C for 18 h after induction with 0.2 mM IPTG (isopropyl β-D-1-thiogalactopyranoside) at an OD_600_ of 0.6-0.8. His-tagged SAM-AMP lyase protein was initially purified by immobilised affinity chromatography (IMAC, 5 mL HisTrap FF column (GE Healthcare)) with gradient elution by increasing the concentration of imidazole from 20 to 500 mM in 50 mM Tris-HCl pH 8.0, 0.5 M NaCl, and 10% glycerol. The untagged protein was recovered by a second IMAC step, after incubation with TEV protease (1 mg TEV protease per 10 mg target protein) at room temperature during overnight dialysis. Size exclusion chromatography (SEC, HiLoad® 16/600 Superdex® 200 prep grade, Cytiva) was conducted for the final purification step in 20 mM Tris-HCl pH 8.0, 250 mM NaCl, 10% glycerol and 1 mM DTT. The homogeneity of SAM-AMP lyase was confirmed by SDS-PAGE (NuPAGE Bis-Tris Gel, Thermo Fisher Scientific), followed by flash freezing and storage at -70 °C.

### Plasmid challenge assay

The method was described previously (22). Briefly, *E. coli* BL21Star cells (Invitrogen) were co-transformed with both pBfrCmr1-6 and pBfrCRISPR_Tet (or pBfrCRISPR_pUC). A single transformed colony was selected for competent cell preparation. Cells were cultivated in LB medium plus antibiotics (100 μg/ml ampicillin and 50 μg/ml spectinomycin) at 37 °C overnight, before 50-fold dilution of the overnight culture into 20 ml LB with the same antibiotics. After OD_600_ reached 0.8-1.0, cell pellets were collected and resuspended in an equal volume of pre-chilled buffer (60 mM CaCl_2_, 25 mM MES, pH 5.8, 5 mM MgCl_2_, 5 mM MnCl_2_). Solutions were incubated on ice for 1 h and subsequently pelleted, before resuspending in 0.1 volume of the same buffer containing 10% glycerol. Competent cells (100 μl) were transformed with 50 ng pRATDuet, pRATDuet-SAM-AMP lyase, pRATDuet-BfrCorA-SAM-AMP lyase or SAM-AMP lyase variants. The transformation mixture was incubated at 37 °C for 2.5 h with the addition of 0.5 ml LB medium, after heat shock at 42 °C for 30 s. A 10-fold serial dilution was applied in duplicate to LB agar plates (supplemented with 100 μg/ml ampicillin and 50 μg/ml spectinomycin) for the determination of the quality of competent cells. The transformants were selected on LB agar containing the same two antibiotics used previously, along with 12.5 μg/ml tetracycline. The full induction was performed in the presence of three antibiotics for selection plus 0.2% (w/v) lactose and 0.2% (w/v) L-arabinose. Plates were incubated at 37 ºC overnight and imaged. The experiment was performed as two independent experiments with two biological replicates and at least two technical replicates.

### Phage lyase prediction, cloning, expression, purification

Phage-encoded SAM-AMP lyase candidates were identified based on sequence similarity to the *C. botulinum* protein. The selected candidate protein (NCBI accession DAN18478.1) is encoded by a *Caudoviricete sp*. phage, identified in the HMP Core Microbiome Sampling Protocol (HMP-A) study of the human viral metagenome (27). Two further close homologues (DAD71688 and DAT17917) were also detected in the dataset.

A codon-optimised synthetic g-block (IDT) encoding DAN18478 was designed and cloned between the *Nco*I and *Bam*HI restriction sites of vector pEhisV5TEV (26) for protein expression. The protein was expressed and purified as described for the *C. botulinum* SAM-AMP lyase, with the addition of a final heparin column (Cytiva) chromatography step, where protein was eluted with a gradient of 10 mM to 1 M NaCl in 20 mM Tris-HCl pH 7.5, 10% glycerol. The purity and identity of the target protein were confirmed by SDS-PAGE and mass spectrometry.

### SEC analysis

To examine the oligomerisation state of *C. botulinum* SAM-AMP lyase wild-type and variants, SEC analysis was performed using a Superose6 Increase 10/300 chromatography column (Cytiva). The column was equilibrated with SEC buffer containing 20 mM Tris-HCl pH 8.0, 250 mM NaCl, 10% glycerol and 1 mM DTT, with a flow rate of 0.5 ml/min. Protein samples were diluted with the same buffer to a concentration of ∼1 mg/ml and 100 μl was injected onto the SEC column. The SEC standards (Bio-Rad) were run under the same conditions, consisting of a mixture of molecular weight markers: thyroglobulin (670,000 Da), γ-globulin (158,000 Da), ovalbumin (44,000 Da), myoglobin (17,000 Da), and vitamin B12 (1,350 Da).

### Generation and purification of SAM-AMP

SAM-AMP was purified for crystallographic experiments using *in vivo* production in *E. coli* followed by high-performance liquid chromatography (HPLC). The *in vivo* production was previously described (22). Briefly, *E. coli* BL21 Star cells containing three plasmids (pBfrCmr1-6, pCRISPR_Tet and pRAT-Duet) were fully induced with 0.2% (w/v) D-lactose and 0.2% (w/v) L-arabinose when the OD_600_ of the 20-fold diluted overnight culture reached between 0.4 and 0.6. The cell culture, mixed with four times the volume of cold phosphate-buffered saline, was subjected to centrifugation at 4,000 x *g* for 10 min at 4 °C. Cells were lysed by resuspending them in a cold solvent mixture (acetonitrile:methanol:water, 2:2:1 by volume), vortexed for 30 s, and stored at -20 °C until needed. After centrifugation at 13,000 x *g* for 10 min at 4 °C, the supernatant was collected, completely dried by evaporation, and resuspended in water for HPLC purification.

SAM-AMP was subsequently purified using a Shimadzu Prominence HPLC system equipped with an HSS T3 HPLC column (Waters 250 mm X 4.6 mm, particle size 5.0 μM). 500 μl samples were analysed by gradient elution with solvent A (10 mM ammonium bicarbonate) and solvent B (acetonitrile plus 0.1% TFA) at a flow rate of 0.3 ml/min as follows: 0-20 min, 0-30% B; 20-20.5 min, 30-100% B. The oven temperature was set at 40 °C and the absorbance was monitored at 260 nm. The peak containing SAM-AMP was collected at a retention time of around 11 min with settings of 5 sec width, 10,000 μV/sec slope and 800,000 μV level. The pure collected samples were dried into powder.

### Crystal structure determination of SAM-AMP lyase

Initial crystallization conditions for SAM-AMP lyase were obtained after sparse matrix screening of the protein assessing 384 crystallisation conditions on a nano-litre scale. SAM-AMP lyase and mother liquor (ML) were mixed in 1:1 or 2:1 protein to ML ratios, in vapour diffusion sitting drop plates. The plates were sealed and left to equilibrate at 293 K. Diffraction quality crystals of wild-type and E71Q variant protein crystallised in the same condition. Following optimisation of the initial hit, the established conditions for crystallisation were 67.6% MPD, 0.1 M HEPES pH 7.35, with a protein concentration of 10 mg mL^−1^. To trap SAM-AMP with the protein, E71Q crystals were soaked with a few grains of SAM-AMP in the original crystallisation drop for 90 min. Prior to data collection all crystals were cryoprotected with ML containing 25% glycerol before cryo-cooling in liquid nitrogen.

X-ray data from WT and E71Q-soaked crystals were collected at a wavelength of 0.9212 Å, at 100 K, on beamline I04-1 at Diamond Light Source. Data were automatically processed using xia2 (28) with DIALS (29) to 1.64 Å (WT) and 1.56 Å (E71Q) resolution, although CC_1/2_ in the outer bin was < 0.5 for both. Consequently, data were truncated at 1.70 Å (WT) and 1.65 Å (E71Q) resolution in AIMLESS (30) prior to structure solution to increase CC_1/2_ above 0.5 in the outer resolution bin for each data set. WT data were phased using PhaserMR (31) in the CCP4 suite (32) utilising a model generated by AlphaFold2 (33) implemented in Colab, with initial B-factors modelled in Phenix (34). The data from E71Q crystals soaked with SAM-AMP were phased using the WT as the model in PhaserMR.

Both models were refined in the same manner, via iterative cycles of REFMAC5 (35) with manual model manipulation in COOT (36). For the ligand bound structure, electron density for MTA-AMP was clearly visible in the maximum likelihood/σA weighted *F*_obs_-*F*_calc_ electron density map at 3σ. The coordinates for MTA-AMP were generated in ChemDraw (Perkin Elmer) and the library was generated using acedrg (37), before fitting of the molecule in COOT. Model quality was monitored throughout using Molprobity (38). Ramachandran statistics for the WT structure are 97.5% favoured, 0% disallowed and for E71Q structure 98.4% favoured, 0.14% disallowed. Data and refinement statistics are shown in Supplementary Table 2. The coordinates and data have been deposited in the Protein Data Bank with deposition codes 9GAD and 9GAB. Sequence alignment of lyases was performed using Clustal Omega (39)and displayed using ESPript3 (40).

### SAM-AMP degradation assay

The activity of SAM-AMP lyase wild-type and variants was determined according to the previously published methods (22). 1 μM SAM-AMP lyase WT and variants were incubated with 100 μM SAM-AMP, S-adenosyl homocysteine-AMP (SAH-AMP) or Sinefungin-AMP (SFG-AMP) generated by *B. fragilis* type IIIB CRISPR complex (22) in 20 mM Tris-HCl pH 7.5, 250 mM NaCl and 0.5 mM EDTA at 37 °C for the indicated time points. The reaction was stopped by adding 2 equivalents of pre-chilled methanol, vortexed for 30 s and centrifuged at 13,000 rpm, 4 °C for 20 min to remove denatured proteins. The supernatant was dried and resuspended in water before HPLC analysis. The activity of the phage-encoded lyase was assessed using the same approach.

### Analytical high performance liquid chromatography

HPLC analyses were conducted as described in (22) using an UltiMate 3000 UHPLC system (Thermo Fisher scientific) coupled with C18 column (Kinetex EVO 2.1 × 50 mm, particle size 2.6 μm) for enzymatic sample analysis.

### Electrophoretic mobility shift assay

1 μM α-^32^P-radiolabelled SAH-AMP, SAM-AMP or Sinefungin-AMP, synthesised using *B. fragilis* Cmr complex as described previously (22), were incubated with varying amounts of purified SAM-AMP lyase (0, 0.1, 0.2, 0.3, 1, 5, 10 μM) in 12.5 mM Tris-HCl pH 8.0, 5% glycerol and 0.5 mM EDTA at 25 °C for 15 min. Reaction samples mixed with ficoll loading buffer were analysed on a native polyacrylamide gel (8 % (w/v) 19:1 acrylamide: bis-acrylamide), with electrophoresis at 200 V for 2 h at room temperature using 1X TBE running buffer. Phosphor imaging was carried out on a Typhoon FLA 7000 imager (GE Healthcare) with photomultiplier tube setting between 700 and 900.

## RESULTS

### C. botulinum SAM-AMP lyase is functional *in vivo*

Some type III-B CorA-containing CRISPR systems replace the *nrn* phosphodiesterase gene with a gene encoding a predicted lyase (Figure 1A) and we previously demonstrated that the *C. botulinum* lyase degraded SAM-AMP into 5’-methylthioadenosine (MTA)-AMP and L-homoserine lactone *in vitro* (22).

**Figure 1.**
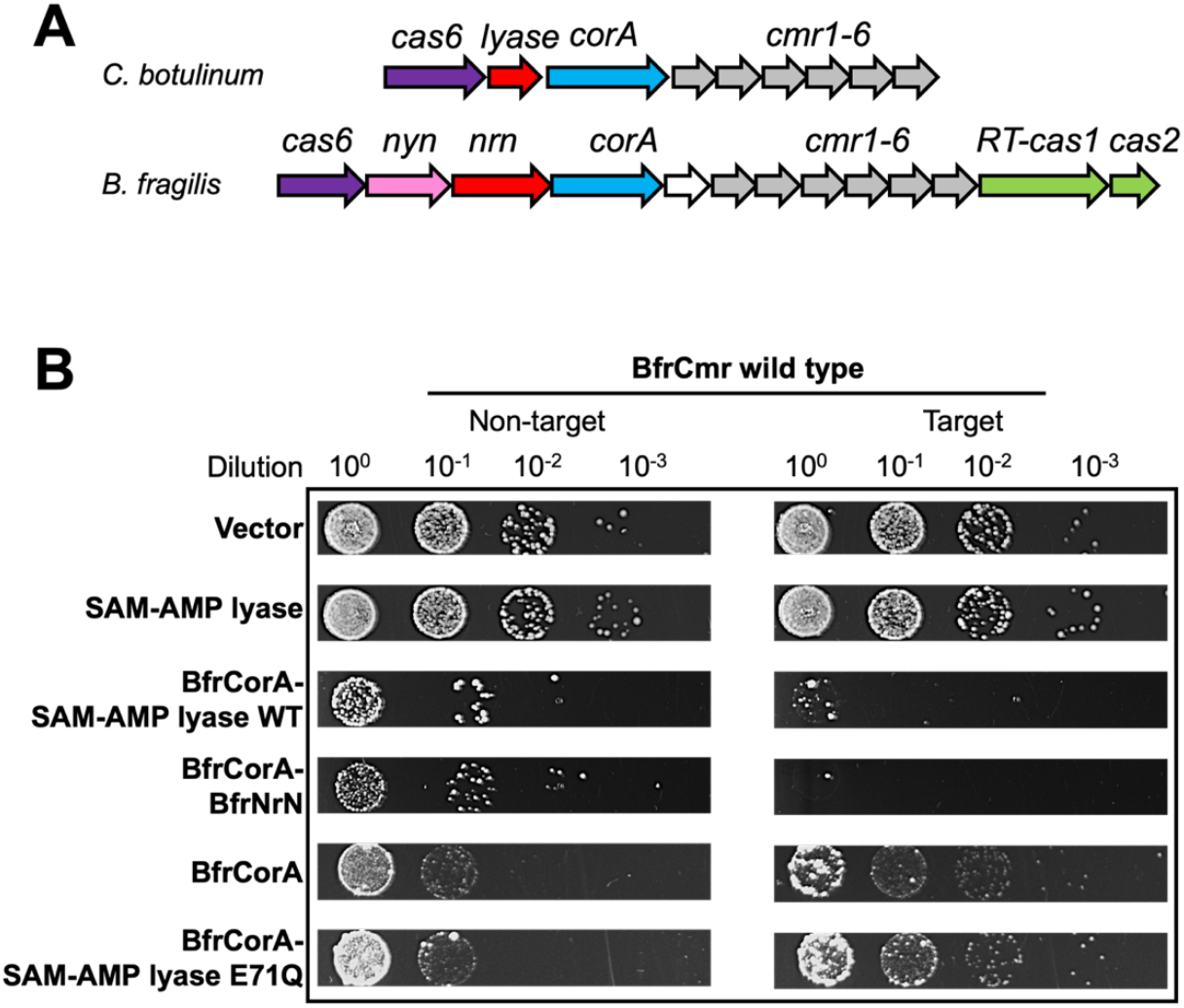
SAM-AMP lyase can replace the NrN phosphodiesterase in a plasmid challenge assay. **A**. Genome context of the CorA-associated type III-B CRISPR systems in *Clostridium botulinum* and *Bacteroides fragilis*. In *C. botulinum*, a predicted SAM-AMP lyase (red) substitutes for NrN (red) in the *B. fragilis* genome, each adjacent to the gene encoding membrane channel protein CorA (blue). The NYN ribonuclease (pink) for crRNA processing (41) and the adaption genes cas1 and cas2 (green) in *B. fragilis* are also shown. The type III-B *cas* genes *cmr1-6* (grey) and *cas6* (purple) are present in both systems. **B**. Plasmid challenge assay. *C. botulinum* SAM-AMP lyase was tested alone or with *B. fragilis* CorA in the context of *B. fragilis* type III-B CRISPR effector (BfrCmr) programmed with target (tetR) or non-target (pUC19) crRNAs. Cells transformed with a pRATDuet plasmid carrying a tetracycline resistance gene served as a vector control. A pRATDuet plasmid expressing both NrN and CorA served as a positive control for plasmid immunity, which was also observed when NrN was replaced with the *C. botulinum* SAM-AMP lyase. Immunity was lost when the predicted catalytic residue was mutated (E71Q) in SAM-AMP lyase.

Given the strict requirement for both SAM-AMP synthesis and degradation in the *B. fragilis* type III-B CRISPR system (22), we investigated whether SAM-AMP lyase could replace the functional role of the NrN SAM-AMP phosphodiesterase. To assess this, we expressed the *B. fragilis* type III-B system in *E. coli* cells, along with a targeting (pCRISPR-Tet) or non-targeting (pCRISPR-pUC) crRNA. Cells were then challenged by transformation with a pRATDuet plasmid harbouring a *tetR* gene together with genes encoding combinations of CorA, NrN and SAM-AMP lyase (Figure 1B). Cells with an active CRISPR system and *tetR* targeting crRNA prevent plasmid transformation (22). As observed previously, CorA did not prevent plasmid transformation unless the NrN protein was also expressed, although some toxicity was observed even for non-targeting crRNA.

Expression of SAM-AMP lyase in the context of an active CRISPR system did not affect transformation efficiency, showing the same transformation level as the vector control. However, co-expression of SAM-AMP lyase and CorA resulted in a reduced number of colonies for the activated CRISPR system, mirroring the phenotype observed for NrN (Figure 1B). Notably, when a SAM-AMP lyase variant with a mutation in the predicted catalytic site (E71Q) was present, colony formation patterns resembled those observed with only CorA expression. These findings confirm the strict requirement for SAM-AMP degradation activity in the CorA-containing system and demonstrate that SAM-AMP lyase functions analogously to the NrN phosphodiesterase in this context, albeit this requirement is still not fully understood.

### Binding and cleavage of SAM-AMP and analogues by SAM-AMP lyase

To assess the substrate specificity of SAM-AMP lyase, we generated unlabelled and ^32^P-radiolabelled versions of SAM-AMP and its analogues SAH-AMP and Sinefungin-AMP, which exhibit structural variability at the sulphur centre of the methionine moiety. HPLC analysis of reaction products demonstrated that neither Sinefungin-AMP nor SAH-AMP were substrates for SAM-AMP lyase (Figure 2B), confirming the predicted requirement of the lyase reaction for a positively charged sulphonium ion. SAM-AMP lyase was incubated with the radiolabelled molecules and subsequently analysed via native gel electrophoresis. SAM-AMP lyase was active in the binding buffer, resulting in a shift in the radioactive signal to a faster migrating band corresponding to MTA-AMP (Figure 2C, top panel). Clear retarded species were observed at higher concentrations of SAM-AMP lyase, which likely represent bound MTA-AMP. In contrast, SAH-AMP and Sinefungin-AMP did not exhibit a shift (Figure 2C), suggesting that these analogues are not tightly bound by SAM-AMP lyase.

**Figure 2.**
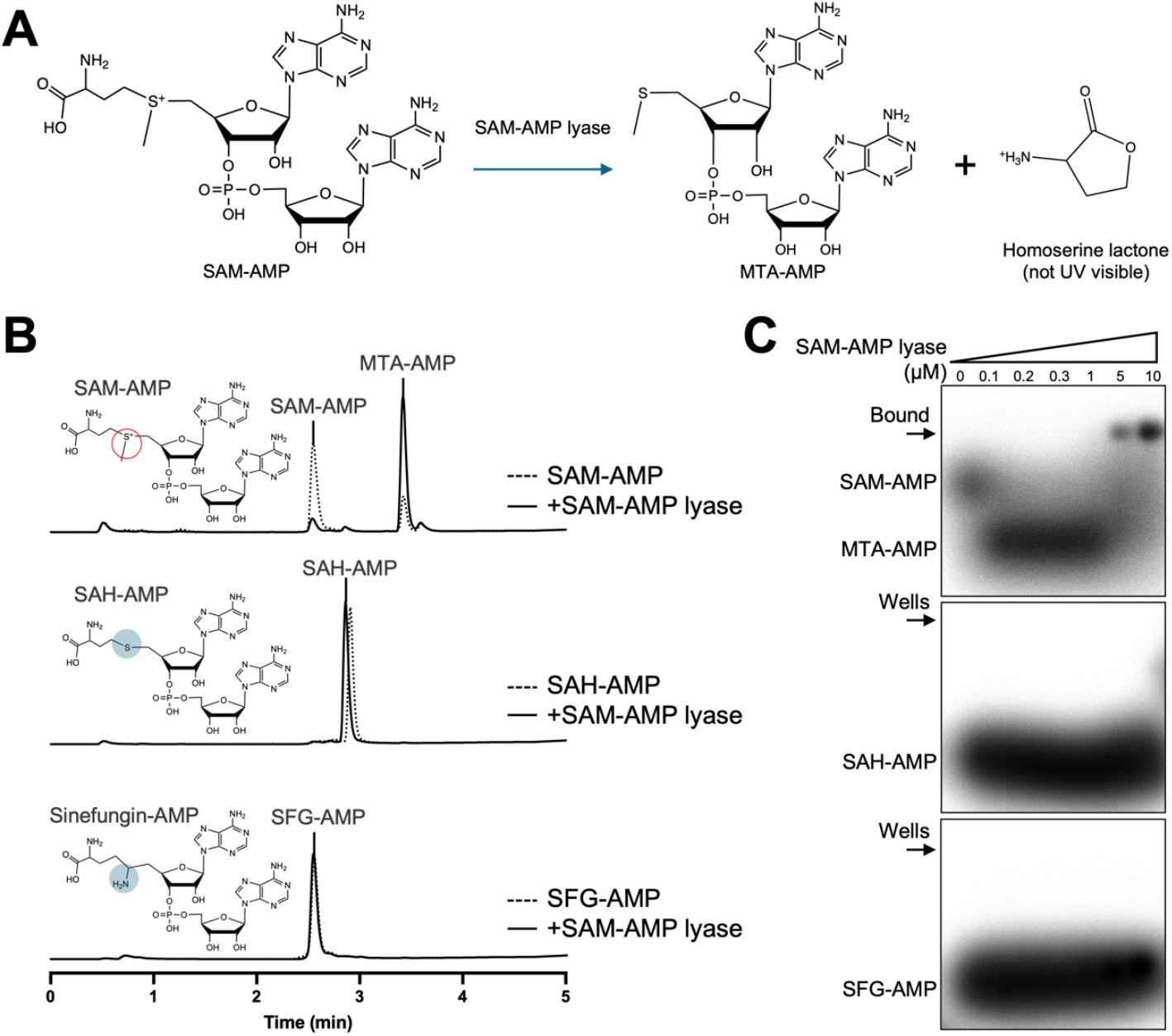
Binding and cleavage of SAM-AMP and analogues. **A**. Schematic representation of SAM-AMP degradation catalysed by *C. botulinum* SAM-AMP lyase. The product homoserine lactone (HL) is not UV detectable. **B**. SAM-AMP lyase specifically cleaves SAM-AMP to MTA-AMP and HL. HPLC analysis of reactions in which SAM-AMP lyase was incubated with SAM-AMP, SAH-AMP or Sinefungin-AMP (SFG-AMP). Circles highlight their structural difference in the sulphur centre. **C**. SAM-AMP lyase binds its product MTA-AMP, but not SAH-AMP or SFG-AMP. 1 μM ^32^P-labelled ligands incubated with 0, 0.1, 0.2, 0.3, 1, 5, 10 μM SAM-AMP lyase were analysed by native gel electrophoresis and imaged by phosphor imaging. Representative figures of three repeats are shown.

### Trimeric structure of *C. botulinum* SAM-AMP lyase

Purified wild-type SAM-AMP lyase (Supplementary Figure 1) was crystallised, X-ray data were collected to 1.70 Å resolution, and the structure was solved using molecular replacement (Figure 3). Crystals of the E71Q variant, targeting a key predicted active site residue, were soaked with SAM-AMP prior to vitrification and data collected to 1.65 Å resolution. The asymmetric unit contains six copies of SAM-AMP lyase, with two trimeric complexes. SAM-AMP lyase exhibits a typical trimeric PII-like protein architecture characterised by a triangular core formed by the four-stranded antiparallel β-sheet of each subunit (Figure 3A). Each monomer possesses a signature ferredoxin fold (β1-α1-β2-β3-α2-β4), featuring a β-sheet on one side and two α-helices on the other. The β2 and β4 strands from each of the three subunits interact in a ‘head-to-tail’ orientation, contributing to most of the trimeric interface. The T-loop connecting β2 and β3, which plays a key role in interactions with target proteins in canonical PII proteins (42,43), is shorter (10-12 residues in SAM-AMP lyase/SAM lyase compared to >20 residues in homologues) and presumably less flexible in SAM-AMP lyase/SAM lyase (given the loops are fully modelled in the lyases, but missing in the homologues, despite similar data resolution). The B-loop, located between α2 and β4, is another conserved loop in the PII protein superfamily which forms interactions with ligands (43).

**Figure 3.**
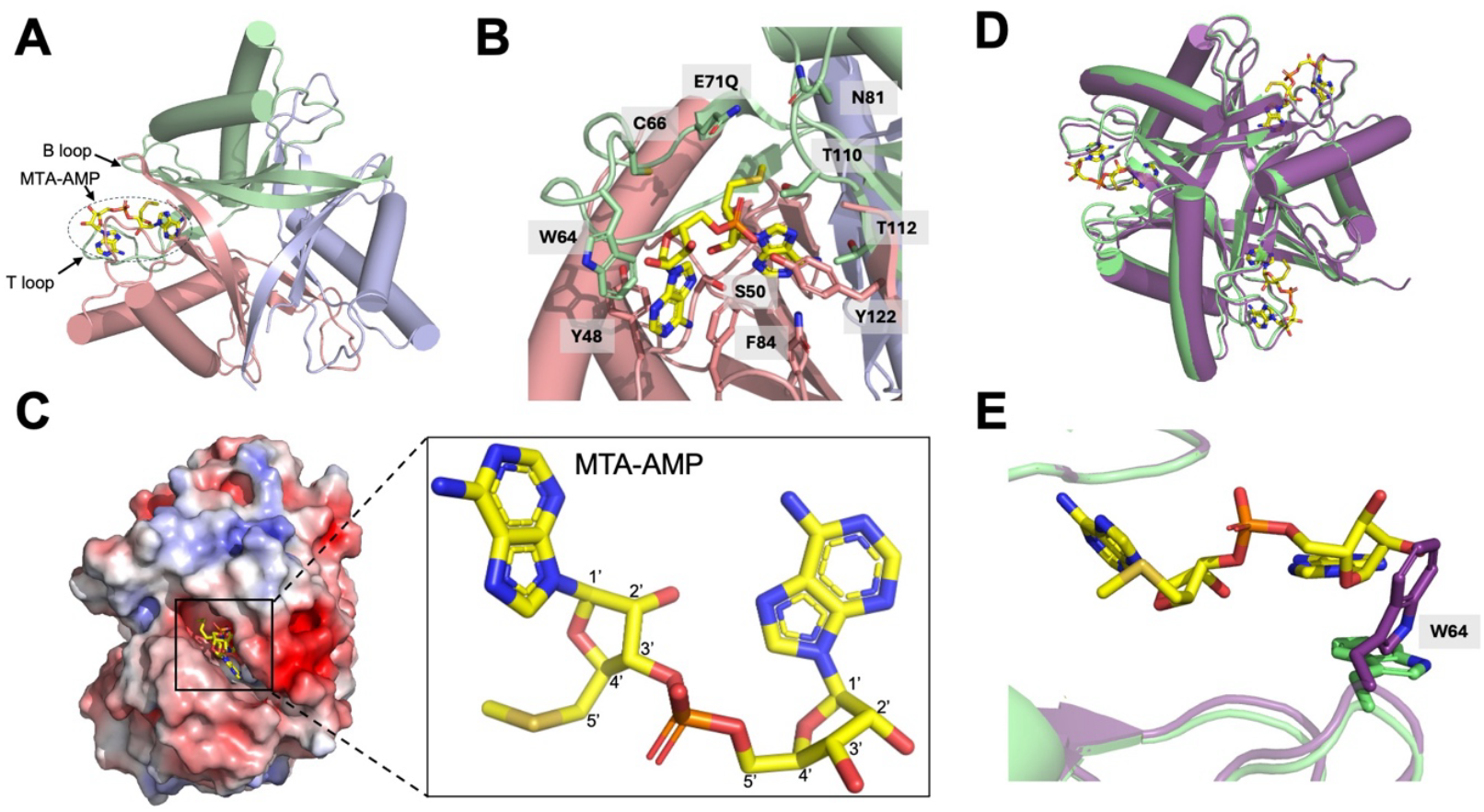
Structure of *C. botulinum* SAM-AMP lyase. **A**. Trimeric structure of SAM-AMP lyase E71Q variant in complex with MTA-AMP. The three monomers are coloured green, blue and salmon. Only one MTA-AMP molecule (yellow sticks), bound at the interface between two subunits, is shown. The T-loop and B-loop in proximity to the MTA-AMP are indicated by arrows. **B**. Binding site for SAM-AMP, with interacting residues shown in the same colours as (A). **C**. Surface representation, coloured by electrostatic potential (blue, (electro)positive; red, (electro)negative; white, non-polar), of SAM-AMP lyase trimer. MTA-AMP molecule is shown as yellow sticks. **D**. Trimeric structure of SAM-AMP lyase E71Q variant (green) in complex with MTA-AMP (yellow sticks) superimposed onto the trimeric structure of wild type apo SAM-AMP lyase (purple). **E**. Binding site for SAM-AMP, showing change in position of Trp64; colours as shown in (D).

### Active site of SAM-AMP lyase

Although SAM-AMP lyase E71Q variant crystals were soaked with SAM-AMP, the product following cleavage, MTA-AMP, was observed in the maximum likelihood/σA weighted *F*_obs_-*F*_calc_ electron density map at 3σ, indicating turnover had occurred. MTA-AMP binds at the interface between two monomers in the trimeric complex. Several residues on the T-loop (comprising residues 59-71) and B-loop (comprising residues 106-111) of SAM-AMP lyase constitute the binding site for MTA-AMP (Figure 3B). The adenine base of AMP is stabilised by π-stacking interactions with the side chains of W64 and F84. T110 and Y122 form hydrogen bonds with the bridging phosphate moiety and the adenine base of MTA interacts with T112 and S50. Other conserved residues Y48, C66, E71, N81, Q108 (Supplementary Figure 2) form the rest of the binding site but make no direct interactions with MTA-AMP (Figure 3B). Note that of the residues discussed, F84, Y122, S50, Y48, and N81 are from a different subunit to the others, demonstrating the significant contributions of neighbouring monomers to ligand binding. The crystal structure of MTA-AMP bound in SAM-AMP lyase confirms the predicted 3’-5’ phosphodiester bond between the SAM and AMP moieties formed by Cas10 (22).

MTA-AMP is buried in a ‘slot-like’, predominantly electronegative, cleft between the subunits of SAM-AMP lyase (Figure 3C), with the predicted position of the carboxyl group in the substrate SAM-AMP orientated towards the B-loop (Figure 3A). Comparison of the apo SAM-AMP lyase and in complex with MTA-AMP reveals almost identical structures (RMSD of 0.3 Å over 123 Cα atoms) (Figure 3D), consistent with the slight conformational change observed in the homologous SAM-lyase (Svi3-3) upon SAM binding (23). This suggests the proteins essentially have a pre-formed binding site for ligands. Given the proximity of the T-loop to the binding site of MTA-AMP, it is perhaps surprising there is no significant movement of this upon binding. However, the side chain of W64, which could only be modelled in some monomers suggesting it is highly flexible, appears to move through 90° from a position which would clash with SAM-AMP to a conformation promoting the π-stacking interaction with the adenine base of the AMP (Figure 3E). This residue may serve as a gate-keeper to the binding site, locking the SAM-AMP substrate in position for catalysis. Interestingly, however, W64 is not conserved in all SAM-AMP lyases (Supplementary Figure 2), so the implication of this movement needs further exploration.

### Comparison of SAM-AMP lyase with SAM lyase (Svi3-3)

Superposition of SAM-AMP lyase in complex with MTA-AMP and SAM lyase in complex with MTA (PDB 6ZM9) gave an RMSD of 2.1 Å over 106 Cα atoms (Figure 4A). Despite the conservation of residues between the two enzymes, there are differences in how the ligands bind (Figure 4B). The adenosine moiety is rotated by approximately 160° in SAM-AMP lyase compared to SAM lyase; whilst the adenine base and methionine moiety for each protein are roughly in the same location of the binding pocket, the ribose sugars are ∼5 Å apart. The structures show E71 of SAM-AMP lyase, positioned at the C-terminus of the T-loop, corresponds to E69 in SAM lyase, which is crucial for catalysis (Figure 4C) (23). The B-loop (residues 106-111 in SAM-AMP lyase) superimposes well with SAM lyase, with the conserved residue Q108 (SAM-AMP lyase) overlapping with Q104 (SAM lyase) (Figure 4C). T112 in SAM-AMP lyase, which interacts with the adenine base, however, is not conserved in SAM lyase (which has an isoleucine in equivalent position) (Figure 4D). The conformation of the T-loop (residues 59-71) differs significantly between SAM-AMP lyase and SAM lyase, meaning residues W64 and C66 in SAM-AMP lyase are not conserved as the equivalent position in SAM lyase is not close enough to MTA to interact directly (Figure 4D). The difference in position is likely due to the addition of the AMP to SAM-AMP substrate, as the loop in SAM lyase would clash with this group. Like SAM-AMP lyase, the interactions made by SAM lyase come from two adjacent monomers in the trimeric protein. Strikingly, however, of the residues from both monomers of SAM-AMP lyase that interact with MTA-AMP, there is only structural conservation of E71 and Q108 in SAM lyase. F84 is conserved by sequence (Supplementary Figure 3), but the phenylalanine in SAM lyase is not observed in the same position structurally (Figure 4E).

**Figure 4.**
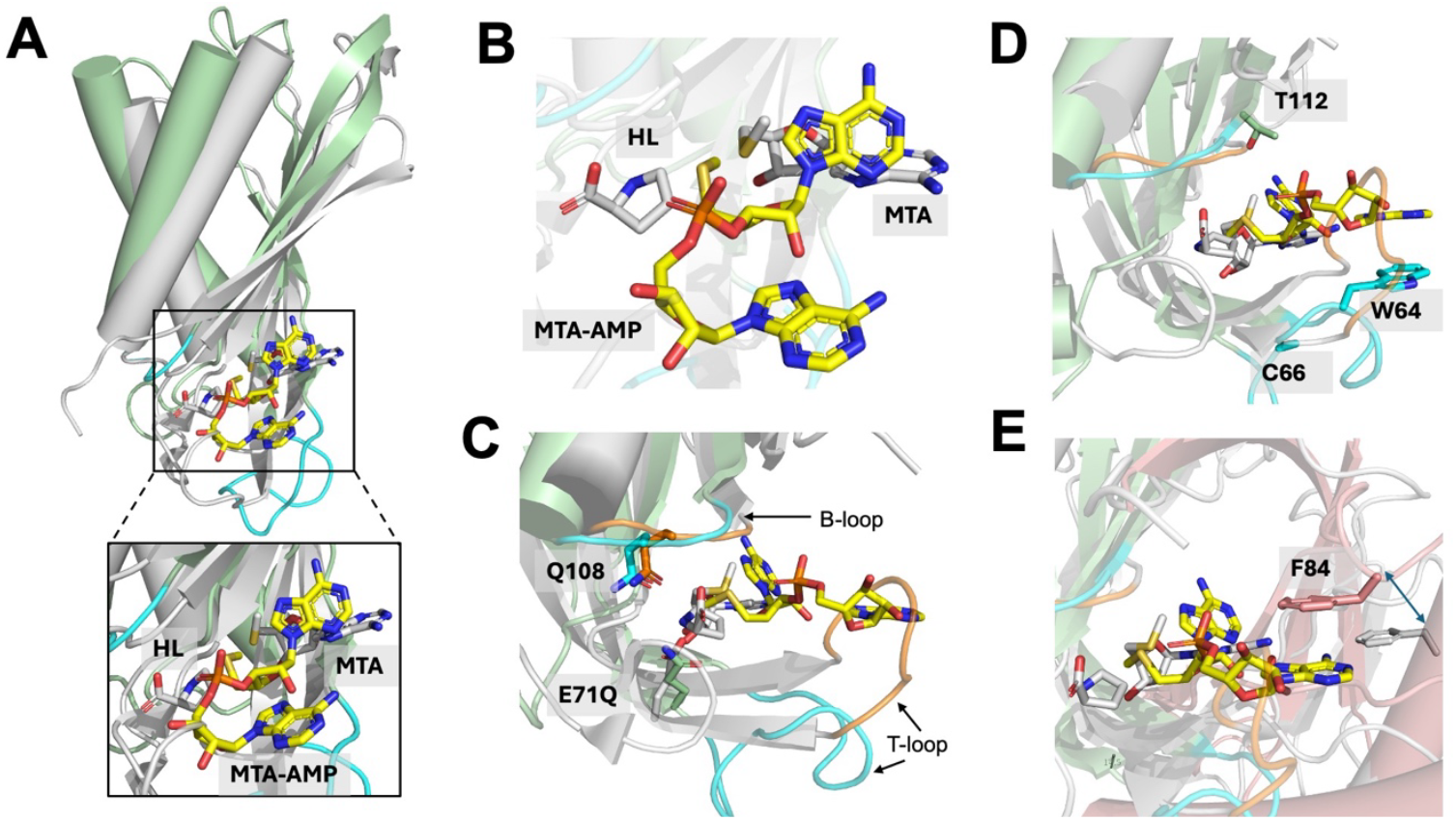
Comparison of structure and active site of SAM-AMP lyase with SAM lyase. **A**. Comparison of monomer structure of SAM-AMP lyase (green) and SAM lyase (Svi3-3, grey) in complex with their ligands, MTA-AMP (yellow sticks) and MTA (grey sticks), respectively. The B-loop (upper) and T-loop (lower) are shown in cyan for SAM-AMP lyase. **B**. Comparison of conformation and position of MTA-AMP (yellow) and MTA (grey) in binding site when the SAM-AMP lyase and SAM lyase are superimposed. **C**. Comparison of active site features for SAM-AMP lyase (green), with the B-loop (upper) and T-loop (lower) shown in cyan in complex with MTA-AMP (yellow), overlapped with SAM lyase (grey), with the B-loop and T-loop shown in orange, in complex with MTA (grey). The only structurally conserved residues between the enzymes are the catalytic residue E71 (shown here as E71Q variant) and Q108. **D**. Comparison of active site features for SAM-AMP lyase and SAM lyase, coloured as in (C), highlighting SAM-AMP residues in the B-loop and T-loop that interact with MTA-AMP, but which are too remote in SAM lyase to form interactions with MTA. **E**. Comparison of active site residues at the dimer interface features for SAM-AMP lyase and SAM lyase, coloured as in (C), with the second monomer shown in salmon for SAM-AMP lyase and grey for SAM lyase. F84, which forms a stacking interaction with the adenine in MTA-AMP, is conserved by sequence with SAM lyase (Supplementary Figure 3), but is not structurally conserved (indicated by arrow), with the equivalent residue in SAM lyase distant from the active site and unlikely to interact with the ligand. All residue numbering shown corresponds to SAM-AMP lyase.

### Probing the role of active site residues

Given the conservation (Supplementary Figure 2) of E71 and Q108 in the SAM-AMP lyases and SAM lyase, which are in the T-loop and B-loop, respectively, we set out to understand if they played a role in substrate binding or catalysis. We also wanted to probe the role played by C66 which is strictly conserved in SAM-AMP lyases. To investigate this the E71Q, C66A, Q108A variants of SAM-AMP lyase were expressed and purified (Supplementary Figure 1). Activity assays revealed that the degradation activity was significantly reduced for the E71Q and Q108A variants, while the C66A variant showed SAM-AMP turnover similar to that of the wild-type. These results suggest significant roles for E71 and Q108 in catalysis, consistent with their conservation between SAM-AMP lyases and SAM lyases, although neither residue is absolutely essential for activity (Figure 5A, B and Supplementary Figure 4A, B).

**Figure 5.**
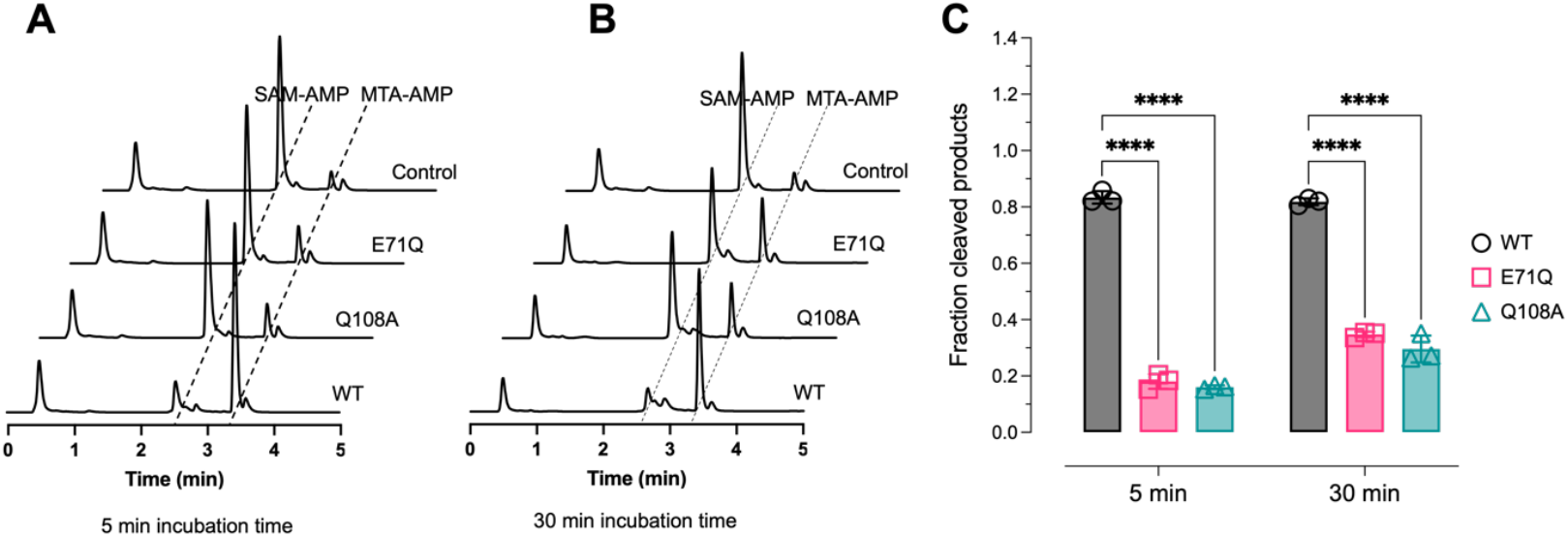
SAM-AMP lyase activity of wild-type and variant proteins. **A**. HPLC analysis of SAM-AMP cleavage activity of wild-type (WT) and variants of *C. botulinum* SAM-AMP lyase, in which 1 μM lyase was incubated with 100 μM SAM-AMP for 5 min (or 30 min, **B). C**. The relative SAM-AMP cleavage activity by quantification of the generation of its product MTA-AMP. Data from three independent experiments are shown, with mean ± SD calculated. Statistical significance was calculated using two-way Analysis of Variance (ANOVA) followed by Dunnett’s multiple comparisons test (\*\*\*\**P*< 0.0001, GraphPad Prism 9).

### Phage encoded SAM-AMP lyases are potential anti-CRISPRs

Nucleotide second messengers such as cOA and SAM-AMP are potent activators of type III CRISPR defence. As such, they are attractive targets for virus-encoded anti-CRISPR (Acr) enzymes that degrade these molecules, neutralising cellular defences (reviewed in (44)). We thus searched for potential phage-encoded SAM-AMP lyases in the NCBI database, identifying a candidate protein (DAN18478) encoded by a phage *Caudoviricetes* species isolate in a viral metagenome dataset isolated from the human microbiome (27). We predict that DAN18478 encodes a trimeric PII-like protein of the SAM lyase family, fused to an N-terminal domain of unknown function (Figure 6A). The gene encoding DAN18478 was recombinantly expressed in *E. coli* and purified to near-homogeneity, allowing biochemical characterisation (Supplementary Figure 5A). The phage protein was an active SAM-AMP lyase, generating MTA-AMP product as observed for the bacterial enzyme, whilst neither SAM nor analogues SAH-AMP or Sinefungin-AMP lyase activity was detected (Figure 6B, Supplementary Figure 5B). Sequence and structural analysis predicted the conservation of conserved acidic residue (E158) at a position equivalent to E71 of SAM-AMP lyase (Figure 6C,D and Supplementary Figure 3). Unfortunately, a E158Q variant of the phage enzyme could not be purified due to low solubility. We were unable to directly test Acr activity *in vivo*, as the *B. fragilis* type III CRISPR complex requires SAM-AMP degradation for immune function in our heterologous *E. coli* experimental system (22). With these caveats, we tentatively assign the name AcrIIIB4 (fourth identified Acr of type IIIB systems) for this phage SAM-AMP lyase.

**Figure 6.**
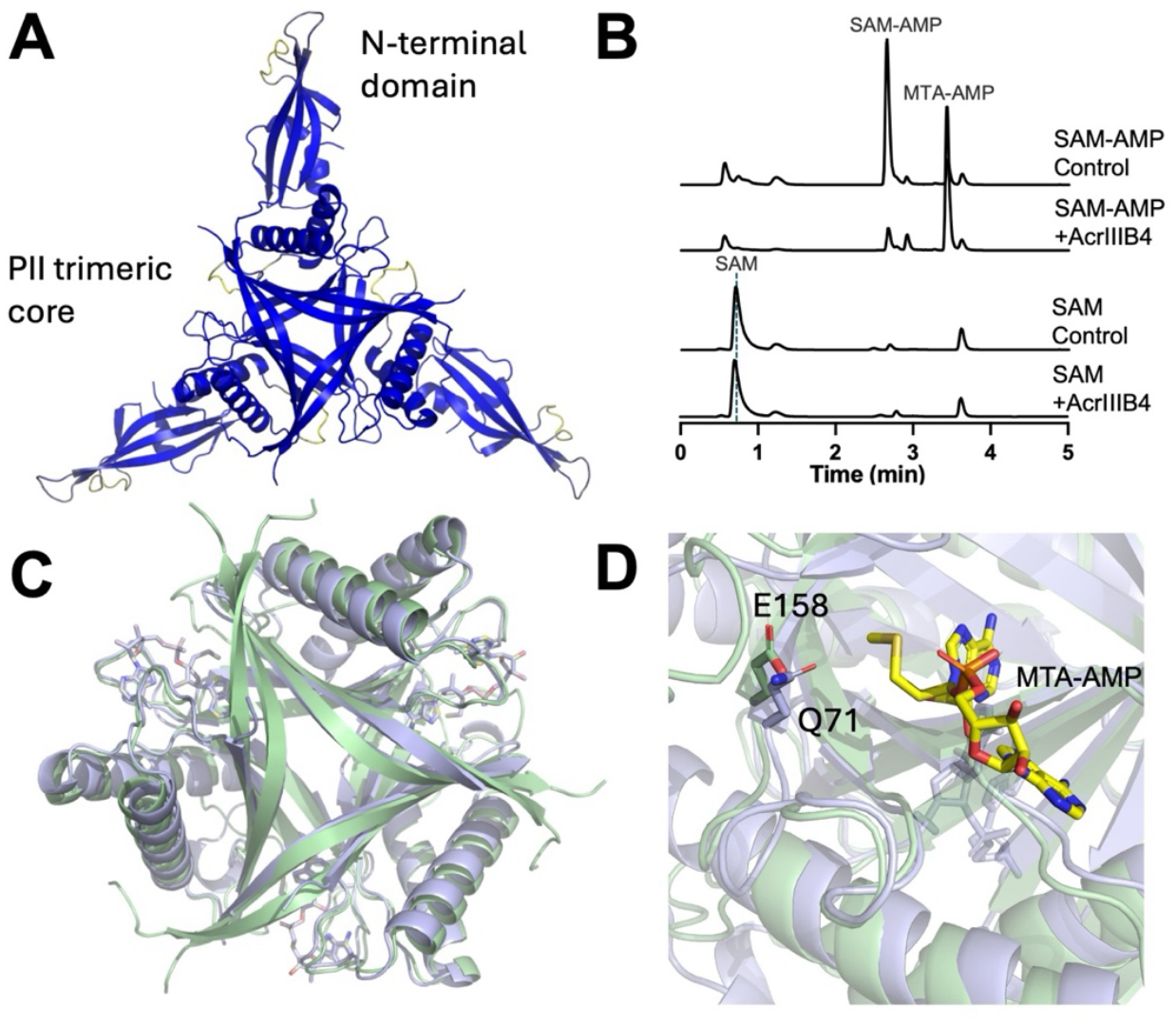
Predicted structure and activity of phage-encoded SAM-AMP lyase AcrIIIB4. **A**. A predicted structure model (AlphaFold3 (45)) of the phage SAM-AMP lyase (AcrIIIB4) coloured according to the pLDDT (predicted local distance difference test) (33). **B**. Specific SAM-AMP cleavage activity of AcrIIIB4. 1 μM AcrIIIB4 was incubated with 100 μM SAM-AMP or SAM for 60 min, followed by HPLC analysis. The cleavage products were MTA-AMP and HL (not UV detectable). **C**. Comparison of predicted AcrIIIB4 trimer model (green) and the CboSAM-AMP lyase E71Q variant with MTA-AMP complex (blue). RMSD is 2.1 Å over 112 Cα atoms. **D**. The ligand binding pocket of AcrIIIB4 (blue) and SAM-AMP lyase (green). The conserved catalytic residues E71 (here mutated to a glutamine) in SAM-AMP lyase and E158 in AcrIIIB4 are shown as sticks. The MTA-AMP reaction product is shown with carbon atoms coloured yellow.

## DISCUSSION

The recent discovery (22) that a clade of type III CRISPR systems respond to infection by synthesising a novel signalling molecule, SAM-AMP, was unexpected. While SAM-AMP binding to a predicted membrane effector of the CorA family is reminiscent of the activation of effectors by cOA species, the observed requirement for SAM-AMP degradation to achieve immunity remains unusual and hard to explain. Although ring nucleases that degrade cOA species are frequently associated with type III CRISPR loci (9,46), they are not essential for immunity and are thought to function in the resetting of the system once an infection has been cleared (47). In contrast, there is no immunity (in a reconstituted, heterologous system) without an enzyme that degrades SAM-AMP (22). Here, we have confirmed that both specialised SAM-AMP NrN phosphodiesterases and lyases can fulfil the same role in the immune response. As these two enzymes generate very different reaction products, this rules out the possibility that a processed form of SAM-AMP is the active signalling molecule. The requirement for this activity remains enigmatic and may require detailed analysis of the mechanism of CorA effector activation and/or studies of cognate host/phage systems.

The structures of SAM-AMP lyase in apo and product-bound forms provide the first detailed view of this enzyme. SAM-AMP lyase is representative of the trimeric PII superfamily structure and closely related to the phage SAM lyase (23) with a key conserved catalytic glutamate residue. For SAM lyase, the suggested mechanism requires dehydration of the carboxylate moiety of SAM in the binding pocket, leading to attack of the sulphonium centre and formation of HL and MTA-AMP in a unimolecular reaction (23). Although SAM-AMP lyase E71Q variant crystals were soaked with SAM-AMP, the electron density revealed only the bound reaction product MTA-AMP, with no density corresponding to HL in the binding pocket. The MTA moiety is bound in a broadly similar position, albeit with variations in conformation and interactions formed, to that observed in the SAM lyase structure. There is an extended binding site for the AMP moiety, which affords SAM-AMP lyase specificity for SAM-AMP over SAM, which is not a substrate. The accommodation of the AMP, however, means a significant difference in the position of the T-loop in the two enzymes, and in fact only two residues are structurally conserved. The co-crystallisation of MTA-AMP provides unambiguous evidence for a 5’-3’ phosphodiester bond linkage, which was previously predicted but could not be proven based on mass spectrometry data alone (22).

The signalling molecules generated by prokaryotic and eukaryotic immune systems represent an attractive target for viral counter-defence measures. Viral “ring nucleases” degrade cA_4_ to neutralise type III CRISPR systems (21), cyclic nucleotides used in CBASS signalling and the cGAMP signalling molecule generated by cGAS in eukaryotic antiviral defence (48-50). Indeed, phage SAM lyase likely functions by disrupting antiviral defence, albeit by targeting a “housekeeping” cofactor (24). The discovery of a phage-encoded enzyme specific for degradation of SAM-AMP fits this paradigm. We envisage that sequestration and degradation of the SAM-AMP molecule prevents activation of the CorA effector and neutralisation of immunity. Although we designate this phage enzyme as the anti- CRISPR AcrIIIB4, functional confirmation *in vivo* will require the study of a cognate host/phage system. Given that the only known function of SAM-AMP is in antiphage defence and that spacers corresponding to phages encoding AcrIIIB4 are found in the genomes of *Leptotrichia* species (27), which utilise CorA-mediated CRISPR defence, this prediction is reasonably well supported.

In conclusion, our work provides the first view of a protein that binds and turns over the recently discovered SAM-AMP signalling molecule in type III CRISPR defence, providing a framework for future studies.

## Data Availability

The protein structure coordinates and data have been deposited in the Protein Data Bank with deposition codes 9GAD and 9GAB.

## Supplementary Data

Supplementary data are available at NAR Online.

## Author Contributions

HC: data acquisition, analysis and interpretation. SM: crystallographic data acquisition, analysis and interpretation. LDP: data acquisition, analysis and interpretation. SG: data acquisition. TM & MFW: funding acquisition, conception, data interpretation. All authors contributed to the drafting and revision of the manuscript.

## Supplementary Figures

**Supplementary Figure 1.**
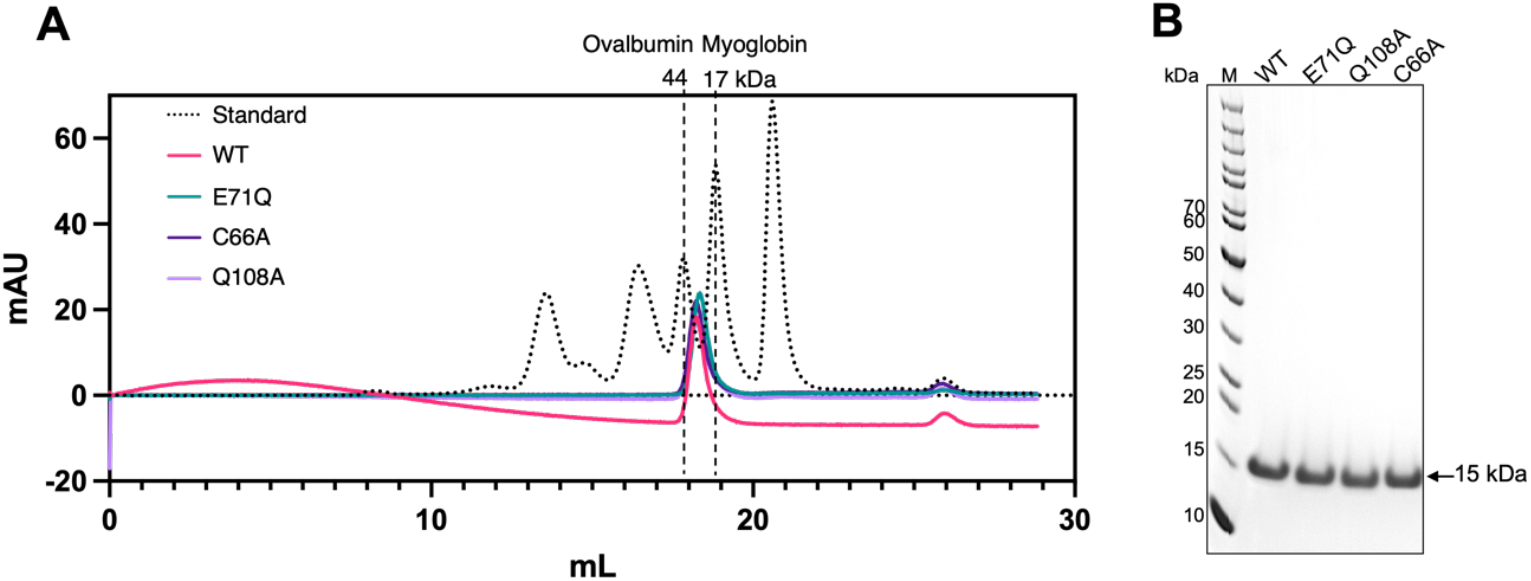
Size exclusion chromatography (SEC) of SAM-AMP lyase wild type and variants. **A**. SEC analysis of SAM-AMP lyase wild type and variants, which all eluted as a trimer (∼30 kDa) between the standards for ovalbumin (44 kDa) and myoglobin (17 kDa). **B**. SDS-PAGE analysis of SAM-AMP lyase wild type and variants. The observed mass of SAM-AMP lyase on the gel is about 15 kDa, consistent with its theoretical monomer mass. M is the molecular weight marker with sizes indicated.

**Supplementary figure 2.**
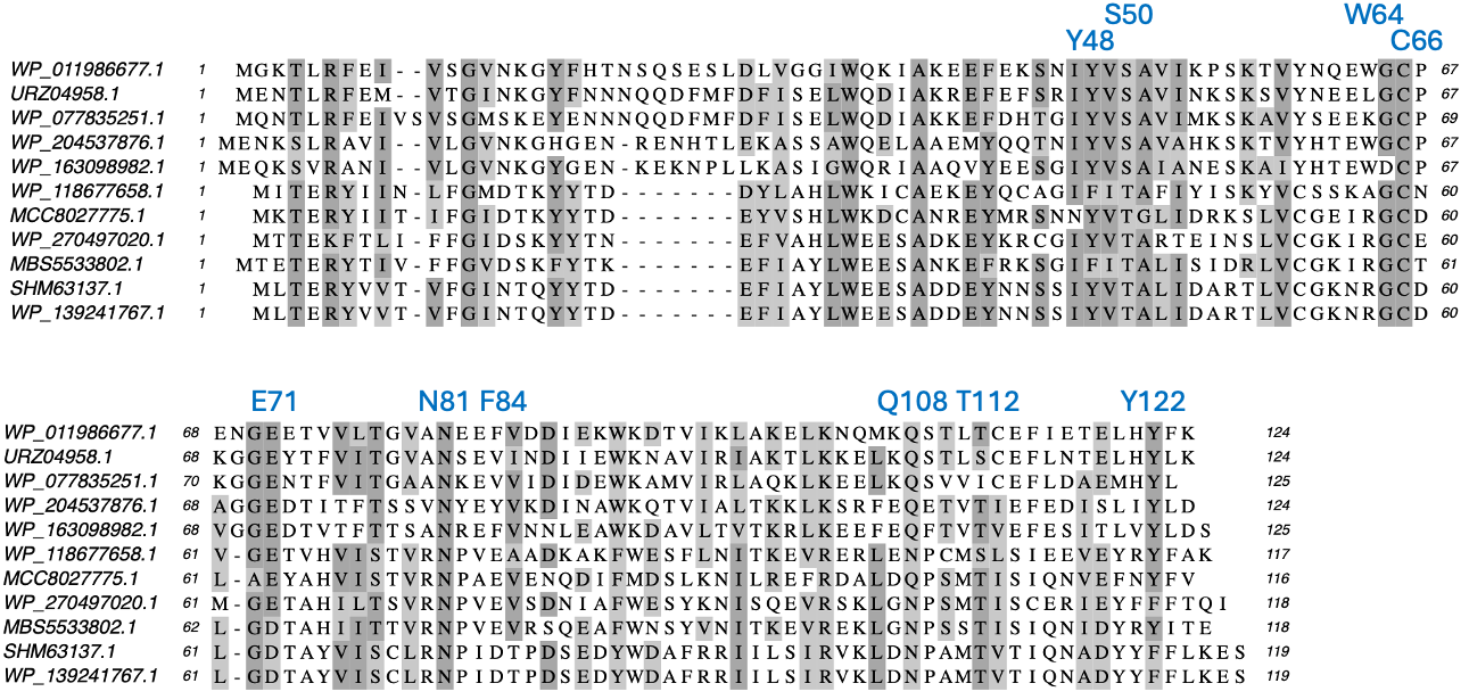
Multiple sequence alignment of a range of bacterial SAM-AMP lyase homologues. Residues mentioned in the text are labelled. NCBI accession codes are shown. The top sequence, WP_011986677.1, is the *C. botunlinum* protein.

**Supplementary Figure 3.**
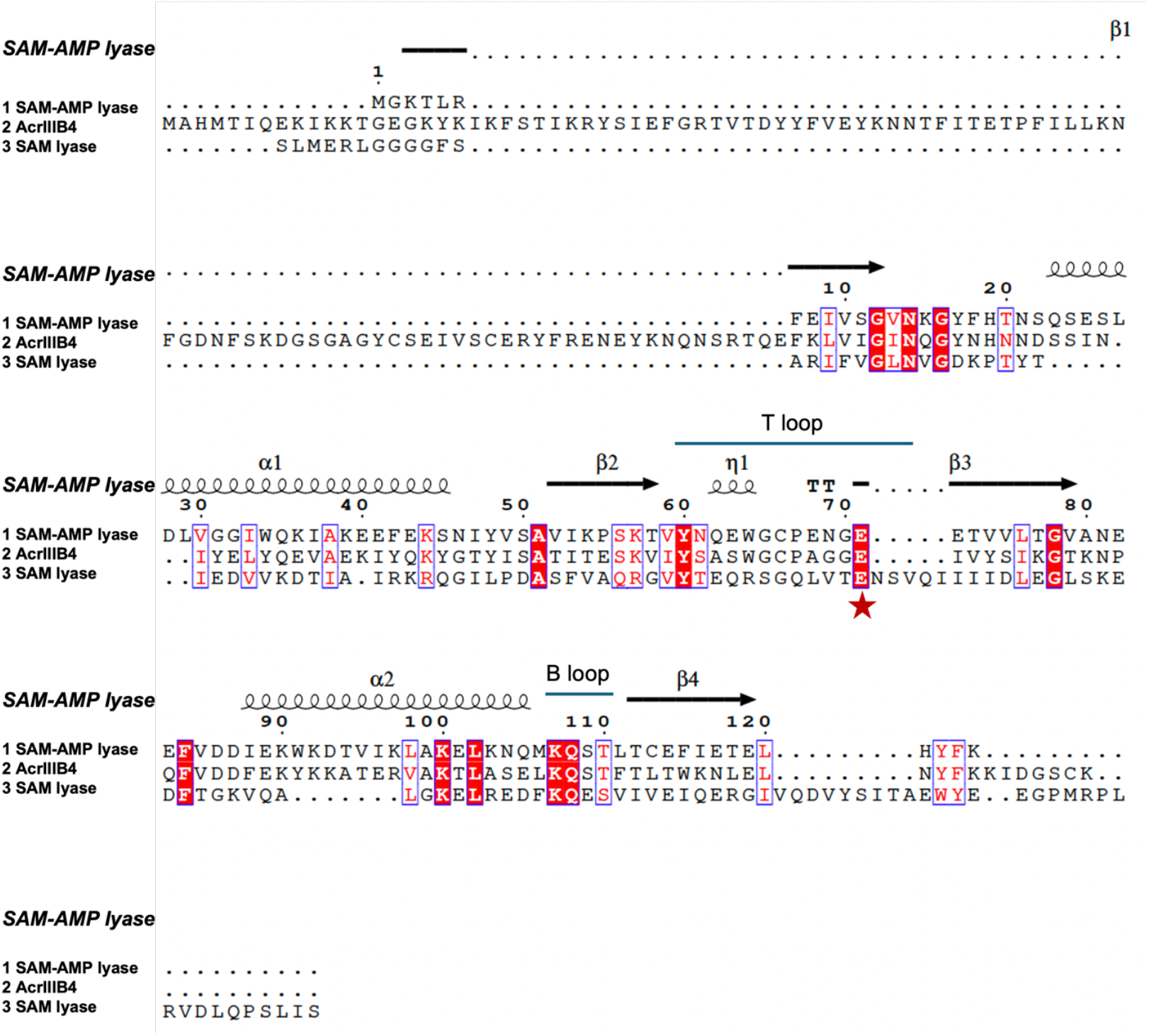
Secondary structure-guided sequence alignment of lyases. Secondary structure of SAM-AMP lyase is displayed above the sequence alignment of SAM-AMP lyase, phage lyase (AcrIIIB4) and SAM lyase (Svi3-3). The white letters on a red background show strictly conserved residues, with the conserved catalytic glutamate residue highlighted as a red star. The red letters with blue frames represent relatively conserved residues. Figure was generated using ESPript3 (40).

**Supplementary Figure 4.**
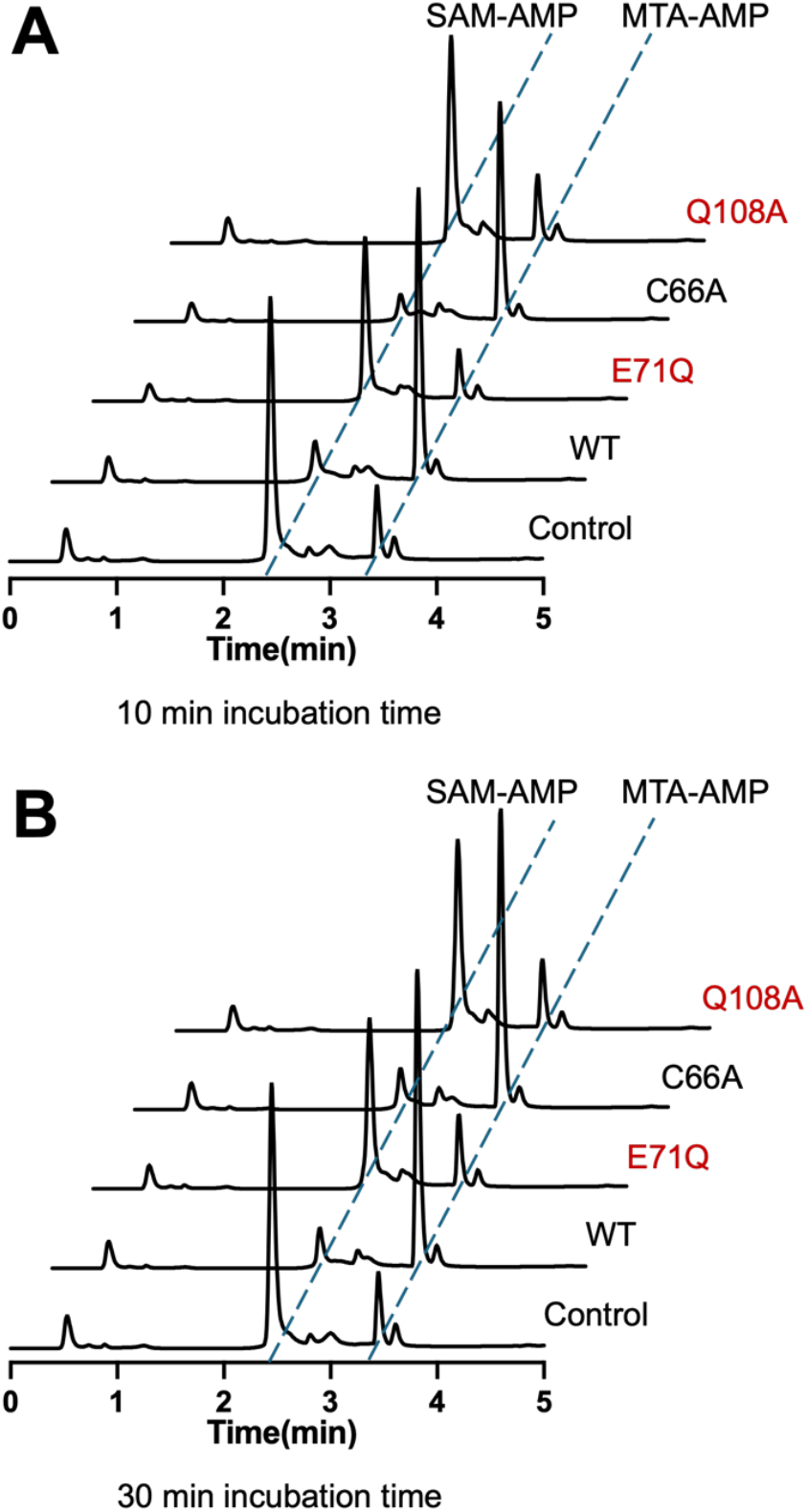
SAM-AMP cleavage activity of SAM-AMP lyase wild type and variants. **A**. Comparison of cleavage activity of SAM-AMP lyase wild type and variants. The generation of cleavage product MTA-AMP was assessed by HPLC. C66A showed similar activity as WT, while the degradation activity was significantly reduced for the variants E71Q and Q108A, after incubation with SAM-AMP for 10 min, or 30 min shown in **B**.

**Supplementary Figure 5.**
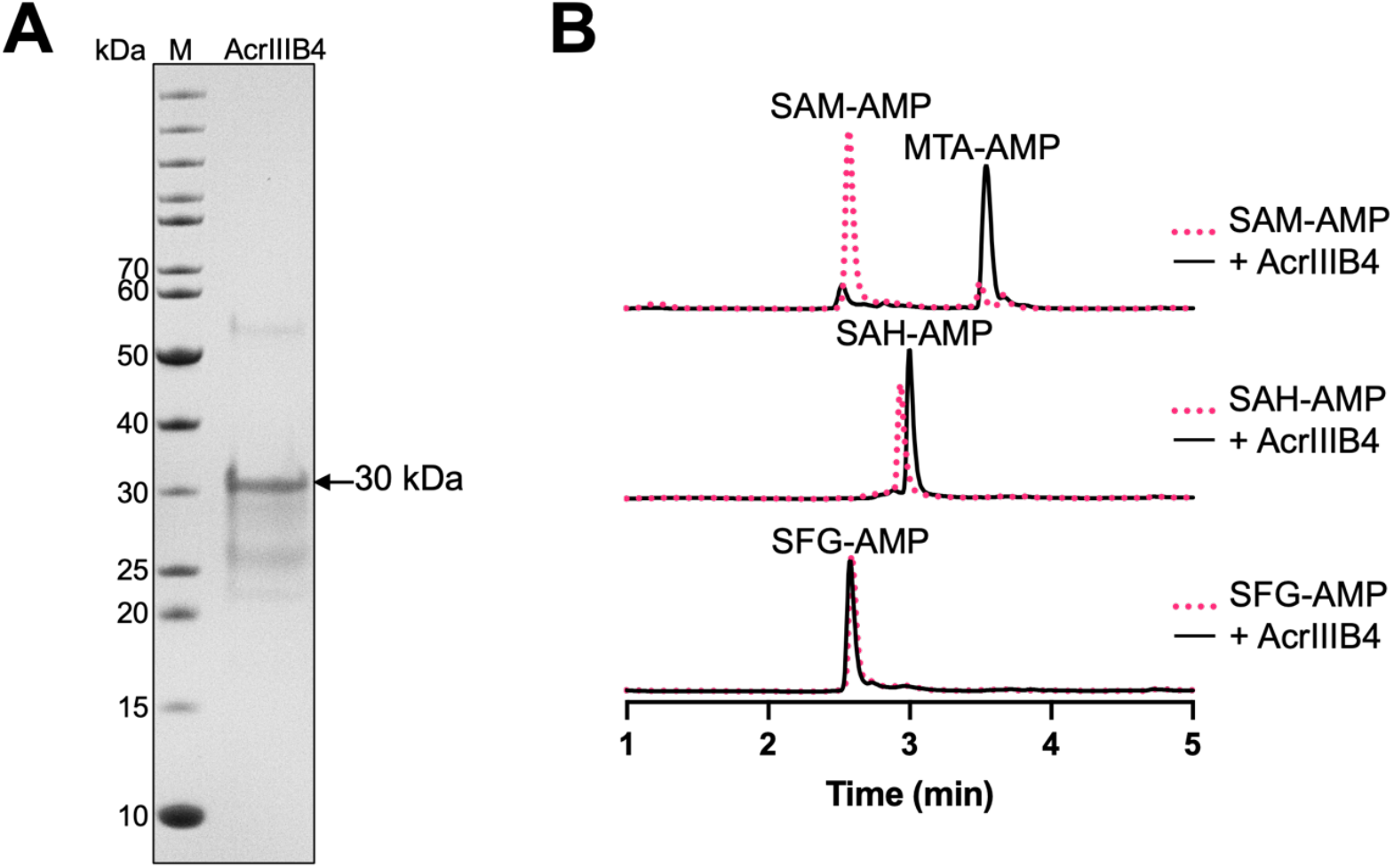
Cleavage of SAM-AMP and analogues by phage SAM-AMP lyase. **A**. SDS-PAGE analysis following recombinant expression of AcrIIIB4. The observed mass of AcrIIIB4 on the gel is about 30 kDa, consistent with its theoretical monomer mass. M is the molecular weight marker. **B**. HPLC traces of AcrIIIB4 with SAM-AMP and analogues SAH-AMP and SFG-AMP. AcrIIIB4 specifically degrades SAM-AMP into MTA-AMP and HL. The traces of control samples are shown in red and AcrIIIB4 reaction samples in black.

**Supplementary Table 1.**
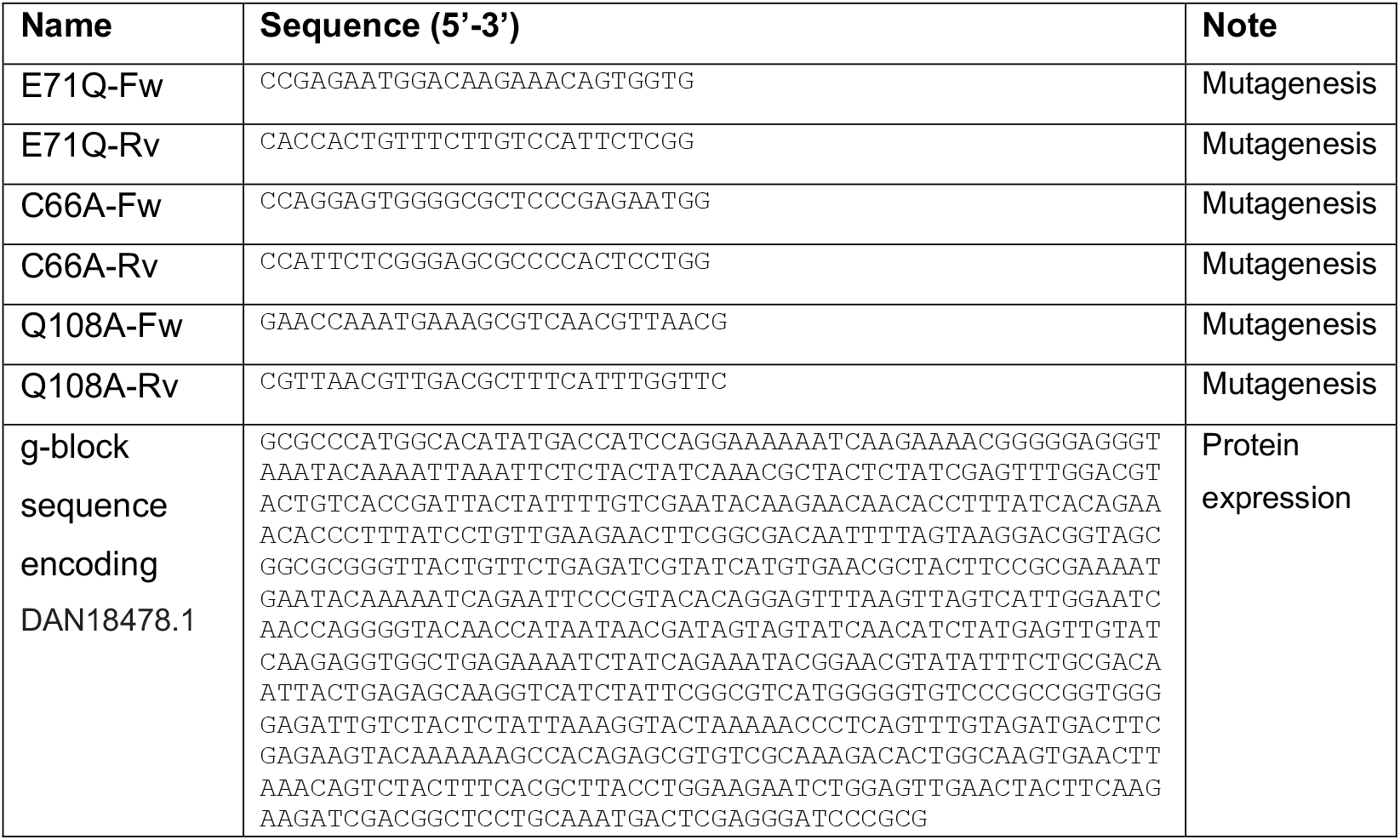
Primers used for SAM-AMP lyase mutagenesis and the synthetic gene encoding AcrIIIB4.

**Supplementary Table 2:** Data processing and refinement statistics for apo WT SAM-AMP lyase and the E71Q variant in complex with MTA-AMP.

|  | WT apo SAM-AMP lyase | SAM-AMP lyase E71Q variant with MTA-AMP |
| --- | --- | --- |
| **Data processing** |  |  |
| Space group | P1 | P1 |
| Cell dimensions |  |  |
| a, b, c (Å) | 54.4, 55.4, 81.7 | 53.8, 55.5, 82.0 |
| $\alpha$, $\beta$, $\gamma$ (°) | 75.7, 72.0, 60.8 | 87.7, 72.6, 61.0 |
| Resolution (Å)<br>(high resolution) | 48.0 – 1.70 (1.73 – 1.70)* | 46.0 – 1.65 (1.68 – 1.65)* |
| $R_{\text{merge}}$ | 0.029 (0.672)* | 0.094 (1.369)* |
| $I/\sigma(I)$ | 13.9 (1.4)* | 10.6 (1.1)* |
| Completeness (%) | 97.6 (96.0)* | 97.5 (95.9)* |
| Average redundancy | 3.6 (3.7) | 7.2 (6.5) |
| CC<sub>1/2</sub> | 0.99 (0.56) | 0.99 (0.63) |
| $V_m$ (Å<sup>3</sup>/Da) | 2.38 | 2.38 |
| Solvent (%) | 48.4 | 48.3 |
| **Refinement** |  |  |
| Unique reflections | 84447 (4431) | 92147 (4548) |
| $R_{\text{work}}$ / $R_{\text{free}}$ | 20.9 / 23.6 | 18.9 / 22.4 |
| Geometric deviations |  |  |
| Bonds (Å) / Angles (°) | 0.008 / 1.55 | 0.008 / 1.52 |
| No. atoms (non H) |  |  |
| Protein | 5971 | 6039 |
| Water | 224 | 514 |
| MTA-AMP | N/A | 252 |
| MPD | 56 | 16 |
| B factors (Å<sup>2</sup>) |  |  |
| Protein | 42.7 | 28.7 |
| Water | 47.3 | 36.6 |
| MTA-AMP | N/A | 20.2 |
| MPD | 43.2 | 49.0 |
| Ramachandran |  |  |
| Favoured / outlier (%) | 97.5 / 0 | 98.2 / 0 |
| Molprobability score / centile (%) | 1.5 / 99 | 0.87 / 100 |

\* Values in parentheses are for the highest-resolution shell.

